# ICM conversion to epiblast by FGF/ERK inhibition is limited in time and requires transcription and protein degradation

**DOI:** 10.1101/157537

**Authors:** Sylvain Bessonnard, Sabrina Coqueran, Sandrine Vandormael-Pournin, Alexandre Dufour, Jérôme Artus, Michel Cohen-Tannoudji

**Affiliations:** Institut Pasteur, CNRS, Unité de Génétique Fonctionnelle de la Souris, UMR 3738, Department of Developmental & Stem Cell Biology, 25 rue du docteur Roux, F-75015 Paris Cedex.; Institut Pasteur, Bioimage Analysis Unit, CNRS UMR 3691, Paris, France.; present address, INSERM UMR935, Paul Brousse Hospital, University Paris Sud, Villejuif, France; present address, Faculty of Medicine, Kremlin-Bicêtre, University Paris Sud, Paris Saclay, France

**Keywords:** inner cell mass, salt and pepper, binary cell fate decision, mouse blastocyst, NANOG, GATA6, protein stability

## Abstract

Abbreviations used in this study:

Epi: epiblast
FGF: fibroblast growth factor
ICM: inner cell mass
PrE: primitive endoderm

**Abstract:** Inner cell Mass (ICM) specification into epiblast (Epi) and primitive endoderm (PrE) is an asynchronous and progressive process taking place between E3.0 to E3.75 under the control of the FGF/ERK signaling pathway. Here, we have analyzed in details the kinetics of specification and found that ICM cell responsiveness to the up and down regulation of FGF signaling activity are temporally distinct. We also showed that PrE progenitors are generated later than Epi progenitors. We further demonstrated that, during this late phase of specification, a 4 hours period of FGF/ERK inhibition prior E3.75 is sufficient to convert ICM cells into Epi. Finally, we showed that ICM conversion into Epi in response to inhibition during this short time window requires both transcription and proteasome degradation. Collectively, our data give new insights into the timing and mechanisms involved in the process of ICM specification.

## Introduction

During early mammalian development, two distinct differentiation steps occur during the formation of the blastocyst. The first one will generate the trophectoderm and the inner cell mass (ICM) followed by the specification of ICM cells into the epiblast (Epi) and the primitive endoderm (PrE). These events are highly coordinated and regulated by a limited number of transcription factors and cell signaling. Epi/PrE formation can be viewed as a three-step model ^1^. First, blastomeres initially co-express the Epi marker NANOG and the PrE marker GATA6 until E3.25 (32-cells) ^2^. Specification of both Epi and PrE is thought to occur asynchronously between E3.25 to E3.75 (64-cells) which is reflected by an ICM composition of cells expressing either NANOG or GATA6 ^3^. These two cell populations ultimately reorganize by a cell sorting process and, by E4.5 (>100 cells), the PrE forms a single cell layer in contact to the blastocoel cavity ^2,4^.

NANOG and GATA6 transcription factors are two key-lineage markers of Epi and PrE formation respectively and have been proposed to mutually repress each other. Indeed, all ICM cells adopt a PrE fate in *Nanog* mutant embryos ^5^ while a reverse situation is observed in *Gata6* mutants ^6,7^.

FGF/ERK signaling pathway is considered as the main regulator of Epi/PrE lineage decision. Genetic inactivation of several members of the FGF pathway including *Grb2* ^3^ and *Fgf4* ^8,9^ impairs PrE formation. Similarly, pharmacological perturbation of FGF/ERK activity also strongly affects PrE/Epi specification ^10,11^. Indeed, when FGFR or MEK is inhibited, all ICM cells adopt an Epi fate. Reciprocally, ICM cells specify into PrE when embryos are cultured in presence of high dose of FGF4. ICM cell plasticity and responsiveness to modulations of FGF activity is progressively lost and around E4.0 cell lineages are determined ^11-13^.

In this study, we have examined in a greater temporal resolution the consequences of modulating FGF/ERK activity during the process of ICM cell specification. We first show that, while being a gradual process, Epi progenitors emerge faster than PrE progenitors. We also refined the windows of ICM cell sensitivity to the modulation of FGF/ERK activity. Particularly, we identified a narrower time window (≤4h) between E3.5 and E3.75 when ICM cells are particularly sensitive to FGF/ERK inhibition. We propose that FGF signaling might act on PrE/Epi specification later than previously thought at the time when Epi progenitors have been formed but PrE progenitors are still emerging. Lastly, we show that both transcriptional and post-translational regulations are required for ICM conversion to Epi upon FGF/ERK inhibition.

## Results

### Sensitivity to FGF/ERK up and down regulation are temporally distinct

Several reports investigated the impact of FGF/ERK signaling modulation on Epi/PrE specification ^10,11,13^. While in absence of FGF activity all ICM cells adopted an Epi fate (NANOG-positive only), a complete reverse situation was observed when FGF signaling was stimulated so that all ICM cells became PrE (GATA6-positive only). Here, we undertook a detailed analysis of the dynamics of ICM cell sensitivity to FGF/ERK modulation during blastocyst formation (i.e from E2.75 to E4.5). To that end, embryos were recovered and cultured from E2.75 to E3.25, E3.25 to E3.75 and E3.75 to E4.5 in presence of either a combination of FGF receptor (PD173074) and MEK (PD0325901) inhibitors or FGF4 ligand. Embryos were next processed to determine by immunofluorescence the proportion of NANOG-positive (Epi progenitors), GATA6-positive (PrE progenitors) and co-expressing ICM cells (Fig. 1). During this time course, we observed the progressive specification of ICM cells as visualized by the reduction of NANOG/GATA6 co-expressing cells (72.4±2.8% at E3.25 (Fig. 1A), 9.3±2.8% at E3.75 (Fig. 1B) and 0% at E4.5 (Fig. 1C)) and confirmed that the salt-and-pepper pattern is mainly established between E3.25 and E3.75 leading to an ICM composition of 40% Epi and 60% PrE at E4.5 as described in ^13^.

**Figure 1.**
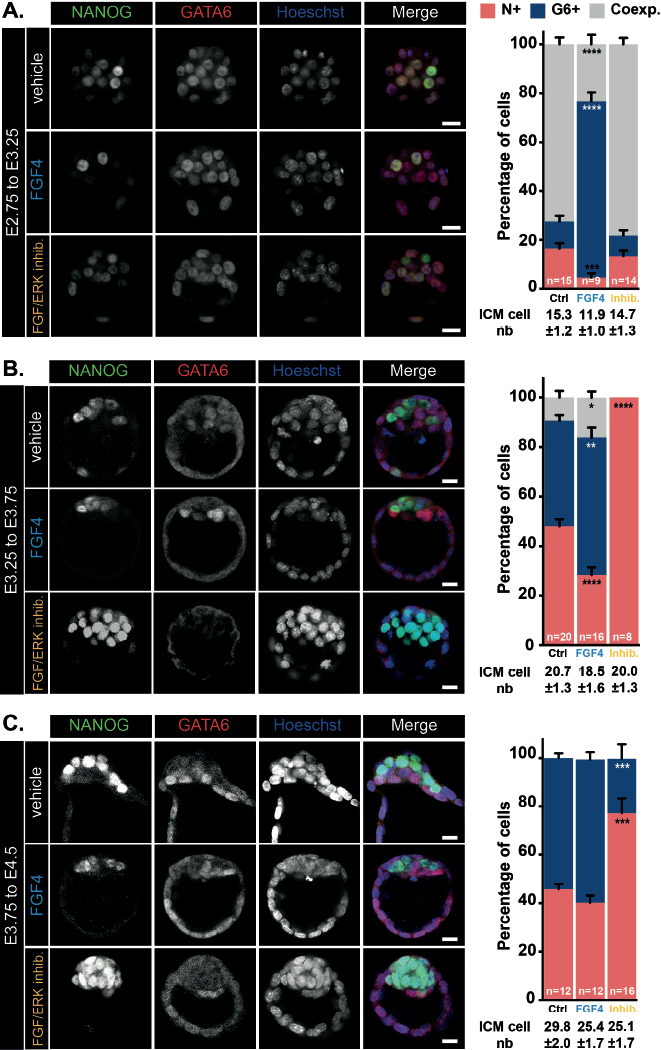
Modulation of FGF/ERK signaling during early mouse development. Immunodetection of NANOG (green) and GATA6 (red) in embryos cultured from E2.75 to E3.25 (**A**), E3.25 to E3.75 (**B**), E3.75 to E4.5 (**C**) and distribution of ICM cells expressing NANOG (N+, red), GATA6 (G6+, blue) or both markers (Coexp., grey). Pictures correspond to a projection of 5 confocal optical slices. Scale bar: 20p.m. Statistical Mann-Whitney tests are indicated when significant (*, p>0.05, **, p < 0.01; ***, p < 0.001, ****, p>0.0001). Error bars indicate SEM. n, number of embryos analyzed.

Upon FGF stimulation, we observed a strong impact on the ICM specification between E2.75 and E3.25 (Fig. 1A) but little (E3.25 to E3.75, Fig. 1B) or no effect (E3.75 to E4.5, Fig. 1C) at later stages. Indeed, at the earliest stages, ICM cells were prematurely specified (72.4±2.8% *vs.* 23.3±3.9% of co-expressing cells in control and FGF4-treated embryos) into PrE progenitors (11±2.3% *vs.* 72±3.7% of GATA6-positive cells in control and FGF4-treated embryos) and at the expense of Epi progenitors (16.5±2.2% vs. 4.7±1.7% of NANOG-positive cells in control and FGF4-treated embryos). We conclude that, as Epi/PrE specification proceeds, ICM cells loose their sensitivity to exogenous FGF stimulation.

On the contrary, FGF/ERK inhibition had no effect between E2.75 and E3.25 (Fig. 1A) but strongly impacted on Epi/PrE specification from E3.25 with a bias towards Epi identity 48±2.7% *vs.* 100% E3.25 to E3.75) and 46±2% vs. 77.2±6.1% E3.75 to E4.5) NANOG-positive cells in control vs. treated embryos). Thus, the highest sensitivity to FGF/ERK inhibition was observed during E3.25 to E3.75 and slightly decreased between E3.75 to E4.5.

Taken together, our data identify two developmental time windows of inner cell sentivity to modulation of FGF/ERK activity: the first one from E2.75 to E3.25 when ICM cells are sensitive to exogenous FGF stimulation but insensitive to FGF inhibition and a second from E3.25 to E4.5 when the inner cells present a reverse sensitivity to the modulation of FGF activity. Thus, ICM cell responsiveness to the up and down regulation of FGF signaling activity are temporally distinct. Interestingly, the period of highest sensitivity to FGF inhibition (E3.25 to E3.75) coincides with the timing of establishment of the salt-and pepper pattern.

### Epi progenitors are specified before PrE progenitors

Having established that, between E3.25 and E3.75, a majority of co-expressing ICM cells progress to a mutually exclusive NANOG/GATA6 expression state, we thought to analyze in greater details the kinetics of emergence of these two, Epi and PrE, progenitor cell populations. We collected E3.25 blastocysts and determined the number of co-expressing NANOG-and GATA6-positive) ICM cells, NANOG-positive Epi progenitors and GATA6-positive PrE progenitors over an 8 hours period of culture (Fig. 2A). Co-expressing ICM cells dropped down from 77% to 10% after 8 hours of culture confirming that most of the ICM cells have been specified during this time window (Fig. 2B–C). Interestingly, during the first 4 hours, the number of Epi progenitors increased rapidly to reach 50% of inner cells. At later time points, no or very little Epi progenitors were produced. Surprisingly, compared to Epi progenitors, production of PrE progenitors was late with a majority of them being produced during the last 4 hours. To confirm our findings, we analyzed the dynamics of SOX17, an additional early PrE marker (Artus et al., 2011; Niakan et al., 2010), during the same timing of culture. We found that the proportion of SOX17-positive, NANOG-negative cells increases mostly during the last 4 hours (Fig. 2D-E). Contrasting with GATA6 stainings, double (SOX17-, NANOG-)-negative ICM cells were observed, which is consistent with our previous observation that *Sox17* shortly follows *Gata6* expression (Artus et al., 2011). Collectively, our data suggest that, while Epi/PrE specification is a gradual process, PrE progenitors tend to be specified later than Epi progenitors.

**Figure 2.**
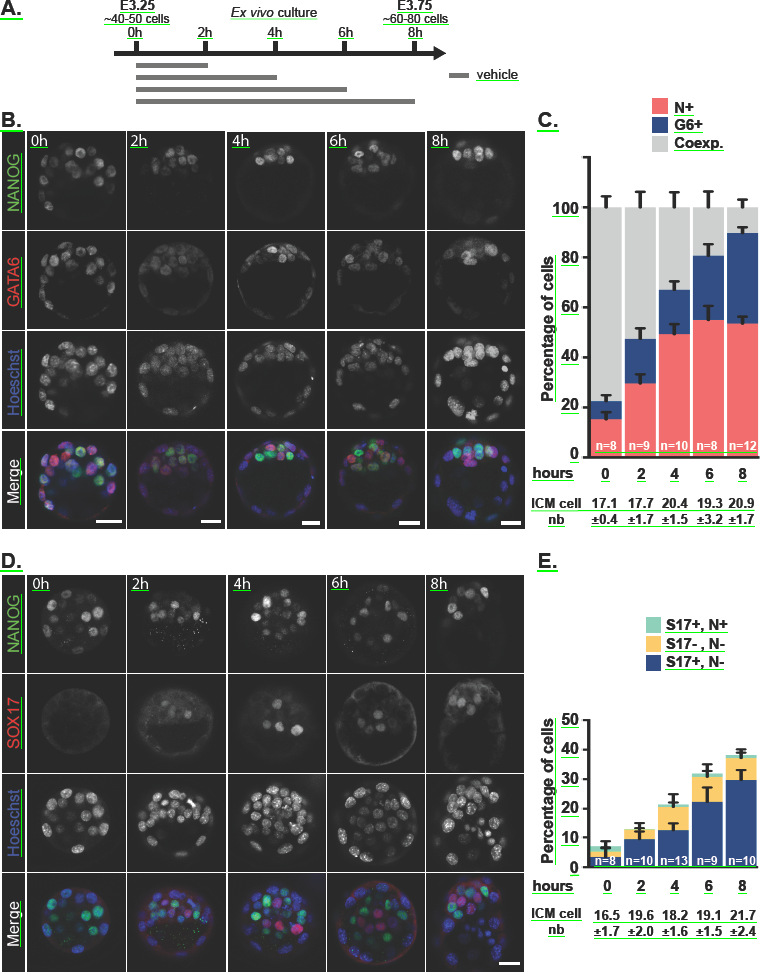
Kinetics of lineage marker expression during blastocyst formation. (**A**) Schematic representation of the culture periods in KSOM+DMSO (vehicle) before analysis. (**B**) Immunodetection of NANOG (green) and GATA6 (red) in embryos cultured for the indicated periods of time. Pictures correspond to a projection of 5 confocal optical slices. (**C**) Distribution of ICM cells expressing NANOG (N+, red), GATA6 (G6+, blue) or both markers (Coexp., grey) in embryos cultured for the indicated period of times. (**D**) Immunodetection of NANOG (green) and SOX17 ( red) in embryos cultured for the indicated periods of time. Pictures correspond to a projection of 5 confocal optical slices. Scale bar: 20μm. (**E**) Number of SOX17+, NANOG-(S17+, N-; blue), SOX17+, NANOG+ (S17+, N+; green) and SOX17-, NANOG-(S17-, N-; yellow) ICM cells in embryos cultured for the indicated period of times. Statistical Mann-Whitney tests are indicated when significant (*, p>0.05). Scale bars: 20pm. (C, F) Error bars indicate SEM. n, number of embryos analyzed.

### ICM cells are sensitive to FGF/ERK modulation during late phase of Epi/PrE cell specification

Our observations that during E3.25 and E3.75, most of ICM cells are specified and exhibit high sensitivity to FGF/ERK inhibitors prompted us to examine in greater details their effect during this time window. We monitored NANOG and GATA6 expression in E3.25 embryos treated for 2, 4, 6 and 8 hours (Fig. 3A). Strikingly, 2, 4 and 6 hours of treatment had no major impact on ICM cell specification (Fig. 3B–C) since their inner cells composition appeared similar to that of control embryos compare Figure 2C and 3C). In particular, GATA6-positive inner cells were still observed in treated embryos. It was only after 8 hours of FGF/ERK signaling inhibition that ICM cells uniformly adopted an Epi fate. Our results suggest that duration of FGF/ERK inhibition might be important to achieve complete ICM conversion to Epi. Alternatively, ICM cells may be sensitive to FGF/ERK inhibition only at a late stage of ICM specification.

**Figure 3.**
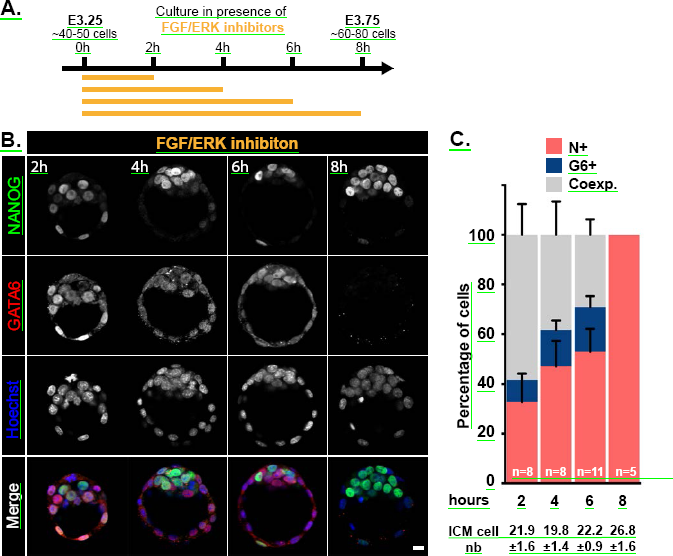
The conversion of ICM into Epi cell lineage requires a 8 hours time period of FGF/ERK inhibition starting from E3.25. (**A**) Schematic of the time schedule of inhibitor treatment. (**B**) Immunodetection of NANOG (green) and GATA6 (red) in embryos cultured for the indicated periods of time. Pictures correspond to a projection of 5 confocal optical slices. Scale bar: 20pm. (**C**) Distribution of ICM cells expressing NANOG (N+, red), GATA6 (G6+, blue) or both markers (Coexp., grey) in embryos cultured for the indicated period of times. Error bars indicate SEM. n, number of embryos analyzed.

To further address these issues, we cultured embryos from E3.25 to E3.75 and added inhibitors during the last 2, 4 or 6 hours of culture (Fig. 4A). Two hours of inhibition were not sufficient to downregulate GATA6 expression in all ICM cells. In contrast, inhibition of FGF/ERK signaling for the last 4 or 6 hours of culture was sufficient to make almost all inner cells expressing only NANOG (Fig. 4B–C). Remarkably, while a 4 hours treatment did not affect ICM specification when applied early (Fig. 3C), it had a major impact when applied late (Fig. 4C). This observation suggests that the effect of FGF/ERK inhibition on ICM cell fate largely depends on the developmental period when inhibitors are applied rather than the duration of the treatment.

**Figure 4.**
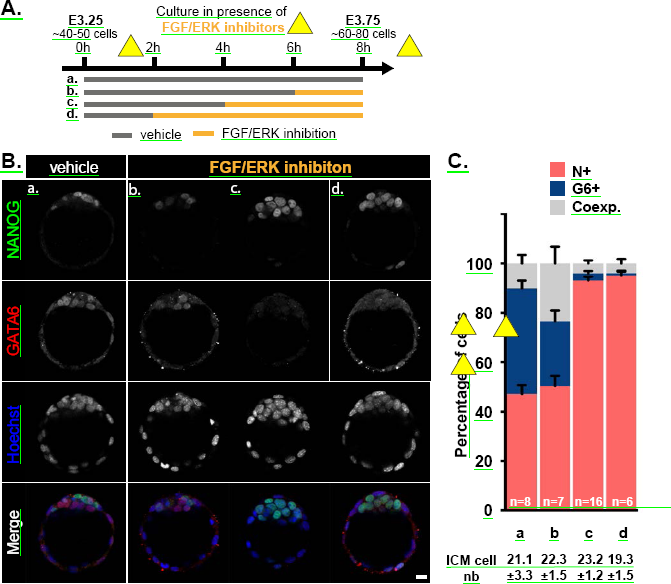
The time window for induced EPI lineage conversion can be narrowed to a 4 hours time period prior E3.75. (**A**) Schematic of the time schedule of inhibitor treatment. Orange and grey lines indicate the culture periods in the presence of inhibitors or DMSO (vehicle), respectively. (**B**) Immunodetection of NANOG (green) and GATA6 (red) in embryos cultured for the indicated periods of time. Pictures correspond to a projection of 5 confocal optical slices. Scale bar: 20p.m. (**C**) Distribution of ICM cells expressing NANOG (N+, red), GATA6 (G6+, blue) or both markers (Coexp., grey) in embryos cultured for the indicated period of times. Error bars indicate SEM. n, number of embryos analyzed.

Even though FGF stimulation had a moderate impact on PrE/Epi cell specification between E3.25 and E3.75 (Fig. 1B), we asked whether ICM cells exhibited differential sensitivity to FGF stimulation (Fig. S1A). We observed a significant bias towards PrE fate at the expense of Epi only when the treatment was applied during the last 4 hours of that period (Fig. S1B-C). Interestingly, only the number of NANOG-only ICM cells was slightly reduced compared to control (Fig. S1Cd) and corresponded to the number of Epi progenitors already present at the onset of FGF4 treatment (Fig. S1Ba). This suggests that FGF4 treatment blocked unspecified ICM cells towards Epi identity rather than affected the fate of already specified Epi progenitors.

Collectively, our data show that, during the time of Epi/PrE specification (i.e between E3.25 and E3.75), the late phase, which corresponds to the time when Epi progenitors have been formed and PrE progenitors are still emerging, is more sensitive to the modulation of FGF/ERK signaling activity than the early phase.

### Mutually exclusive NANOG/GATA6 expression can occur in absence of transcription or proteasome degradation

Our data show that ICM cells can be instructed towards an Epi fate in a narrow time window 4 hours or less) implying rapid and dynamic regulation of GATA6 and NANOG levels. We thus analyzed the contribution of transcription and proteasome degradation on ICM cell specification during that period with and without FGF/ERK inhibition. Embryos were cultured from E3.25 to E3.75 and treated with flavopiridol, a potent RNA Polymerase II inhibitor ^14^ or with the proteasome inhibitor MG132. Treatment was initiated one hour before the addition of FGF and MEK inhibitors and maintained until the end of the culture (Fig. 5A and Fig. S2). We first confirmed that the 1h pretreatment with flavopiridol (but not with MG132) inhibited transcription as visualized by the reduction of *Rps17* pre-mRNA ( Fig. S2A) and did not affect NANOG and GATA6 expression in pretreated ICM cells (Fig. S2B). After 5 hours, flavopiridol treatment lead to a marked reduction of both *Rps17* pre-and mature mRNA while MG132 treatment affected the level of *Rps17* pre-mRNA only.

**Figure 5.**
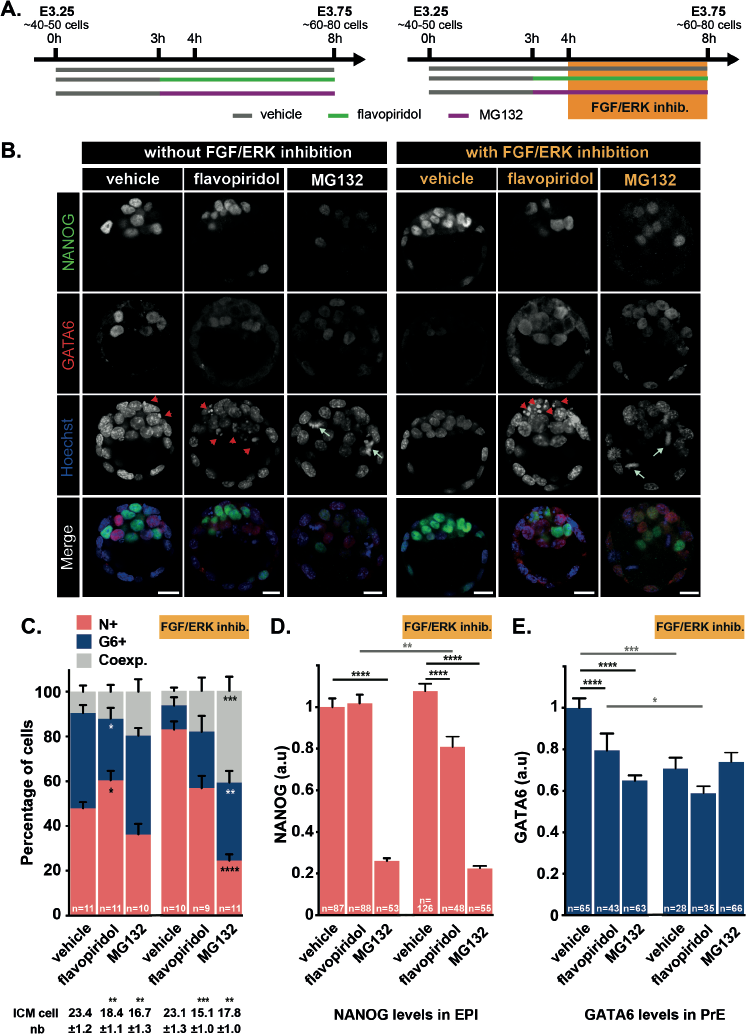
Effect of modulating transcription and proteasome activity during ICM to EPI conversion. (**A**) Schematic of the time schedule of inhibitor treatment. Orange box indicates the 4 hours treatment with FGF/ERK inhibitors prior E3.75. Green, purple and grey lines indicate the culture periods in the presence of flavopiridol, MG132 and DMSO (vehicle), respectively. (**B**) Immunodetection of NANOG (green) and GATA6 (red) in embryos cultured in presence/absence drug treatment. Pictures correspond to a projection of 5 confocal optical slices. Scale bar: 20μm. Red arrowheads: pyknotic nuclei; light green arrows: metaphase. (**C**) Distribution of ICM cells expressing NANOG (N+, red), GATA6 (G6+, blue) or both markers (Coexp., grey) in cultured embryos. Error bars indicate SEM. n, number of embryos analyzed. (**D**) Quantification of NANOG levels in Epi cells (NANOG-positive). (**E**) Quantification of GATA6 levels in PrE cells (GATA6-positive). Error bars indicate SEM. n, number of cells analyzed. Statistical Mann-Whitney tests are indicated when significant (*, p < 0.05; **, p < 0.01; ***, p < 0.001, ****, p>0.0001).

Next, we monitored the consequences of flavopiridol or MG132 treatment on embryonic development. In the presence of flavopiridol, embryos showed a clear reduction of ICM cell number (Fig. 5C) as a probable consequence of increased cell death (Fig. 5B, arrowheads). MG132-treated embryos also exhibited a reduction of ICM cell number (Fig. 5C) likely caused by a block in mitotic progression. Indeed, MG132 treated embryos exhibited a high number of metaphases (Fig. 5B, arrows) reflecting the known function of the anaphase-promoting complex in targeting mitotic proteins for degradation by the proteasome ^15^. Inhibition of neither RNA polymerase II or proteasome degradation prevented the establishment of NANOG and GATA6 exclusive expression by inner cells (Fig. 5C), suggesting that the treatment did not dramatically interfere with ICM specification.

We observed a slight increase of the proportion of Epi cells at the expense of PrE cells in presence of flavopiridol. Interestingly, flavopiridol treatment resulted in a 20% decrease in GATA6 levels in PrE progenitors while NANOG levels in Epi progenitors remained unchanged (Fig. 5D-E). This is consistent with previous reports showing that a significant but moderate reduction of GATA6 in PrE cells in *Gata6+^/-^* embryos leads to an imbalance PrE/Epi specification ^6,7^. Together, our data show that acquisition of PrE fate critically requires active transcription during the late phase of specification.

In contrast, MG132 treatment did not affect the NANOG+/GATA6+ inner cells ratio. Nonetheless, strong reduction in GATA6 levels (40%) in PrE progenitors and in NANOG levels (80%) in Epi progenitors were observed in MG132-treated embryos (Fig. 5B, D-E). Thus, our data show that proteasome activity positively regulates the levels of both transcription factors during that period pointing to additional regulatory mechanisms that remain to be discovered.

We also noticed the presence of GATA6-positive cells in the trophectoderm of MG132-treated embryos, a situation which was rarely observed in control or flavopiridol-treated embryos at late blastocyst stage (Fig. 5B). This observation suggests that GATA6, but not NANOG, levels are negatively regulated by the proteasome in the trophectoderm cell lineage.

### ICM conversion to Epi following FGF/ERK inhibition requires both transcription and proteasome activity

We next analyzed the role of transcription and proteasome degradation on the action of FGF and MEK inhibitors on Epi/PrE specification. In this paradigm, treated embryos are thought to up-regulate NANOG levels and down-regulate GATA6 so that, *in fine,* all ICM cells become exclusively NANOG-positive. Embryos were pretreated for 1h with flavopiridol or MG132 before addition of the inhibitors to the culture medium for 4 additional hours (Fig. 5A). Interestingly, both treatments impaired the effect of FGF/ERK inhibition. Indeed, while control embryos incubated with FGF/ERK inhibitors contained almost exclusively NANOG-positive inner cells, flavopiridol-and MG132-treated embryos contained GATA6 expressing inner cells and essentially resemble to embryos treated with drugs but no inhibitors Fig. 5B–C). The higher proportion of doublepositive cells in embryos treated with MG132 and FGF/ERK inhibitors compared to MG132 alone (40.8±6.3 *vs* 19.7±5.5, p<0.005, Fig. 5C) may be due to the upregulation of NANOG expression in PrE progenitors upon FGF/ERK inhibition together with incomplete downregulation of GATA6 in absence of proteasome activity.

Consistent with the role of FGF/ERK signaling on GATA6 expression ^3,8^, we found reduced GATA6 levels in PrE cells from embryos treated with FGF/ERK inhibitors (Fig. 5E). Absence of further reduction in presence of flavopiridol or MG132 suggests that FGF/ERK regulates GATA6 levels at both transcriptional and posttranscriptional levels. It has been previously reported that FGF/ERK inhibition leads to marked upregulation in NANOG levels in Epi of E4.5 (>100 cells) embryos ^7^. In E3.75 embryos treated with FGF/ERK inhibitors, we found no change in NANOG levels (Fig. 5D) indicating that ICM conversion to Epi does not require deregulated NANOG levels and that FGF/ERK signaling likely controls NANOG levels in Epi after specification. In ES cells, FGF/ERK signaling has been shown to directly repress *Nanog* transcription ^16^. During specification of ICM cells, the link between FGF/ERK signaling and *Nanog* transcription is likely different since NANOG levels were reduced in Epi progenitors of embryos treated with FGF/ERK inhibitors and flavopiridol but not with flavopiridol alone (Fig. 5D).

Collectively, our data show that FGF/ERK inhibitor activity on ICM cell conversion is both dependent on transcription and proteasome degradation.

## Discussion

In this study, we investigated the timing of ICM cell specification into Epi and PrE cell fate and observed that while being a gradual process, the specification of Epi progenitors precedes PrE progenitors (Fig. 6). This is maybe not surprising since PrE specification depends on FGF4 ligand, which is assumed to be secreted by Epi cells once specified ^17^. Importantly, our study redefines the windows of competence during which ICM cells can respond to experimental modulation of FGF/ERK signaling activity. Lastly, we propose that the effect of FGF/ERK inhibition on ICM cells requires transcription and protein degradation.

**Figure 6.**
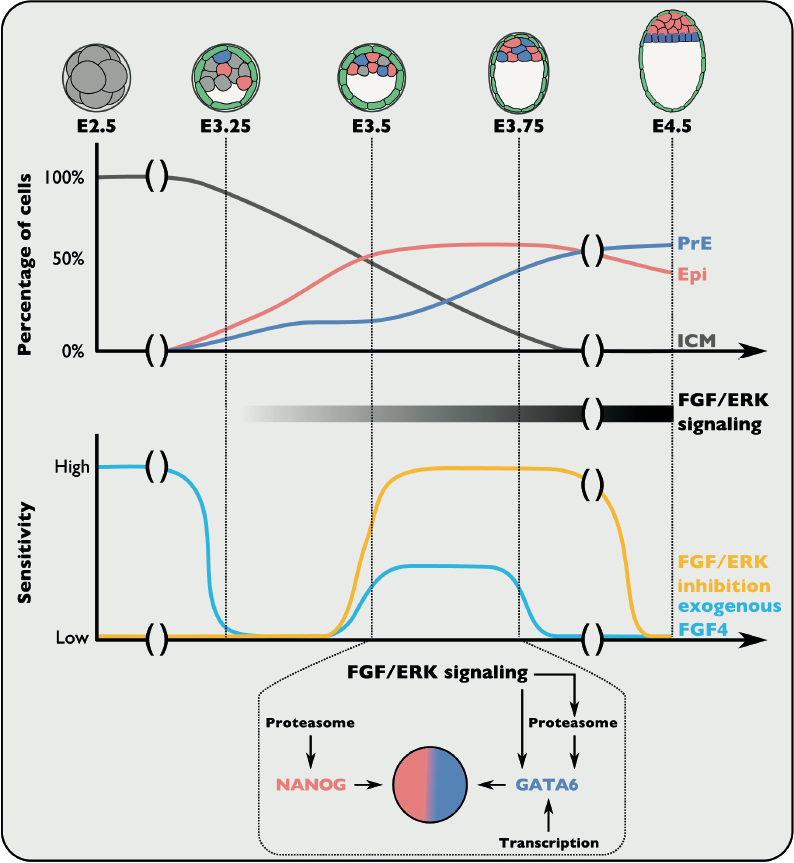
Model of temporal dynamics of ICM cell specification. Specification into Epi (red) or PrE (blue) is a progressive and asynchronous process that occurs for a majority of ICM cells (grey) between E3.25 and E3.75. The formation of Epi progenitors precedes that of PrE progenitors. ICM cell responsiveness to the modulation of FGF/ERK signaling varies over time. First, between E2.5 and E3.25, exogenous FGF4 treatment efficiently diverts unspecified ICM cell from Epi fate. Then, between E3.25 and E3.5, ICM cells are globally insensitive to modulation of FGF/ERK signaling. Between E3.5 and E3.75, remaining unspecified ICM cells but not already specified Epi progenitors are able to respond to exogenous FGF4 leading to a moderate shift in Epi/PrE specification upon treatment. FGF signaling and ERK phosphorylation increases in ICM cells and PrE progenitors during this time window. Accordingly, embryos become highly sensitive to FGF/ERK inhibition leading to the complete ICM conversion to Epi. During that period, proteasome degradation and transcription control NANOG and GATA6 levels in Epi and PrE progenitors. Downregulation of GATA6 levels in PrE progenitors upon FGF/ERK inhibition is partially mediated by the proteasome. After E3.75, responsiveness to exogenous FGF4 is lost when all ICM cells become specified while PrE progenitors are still able to respond to FGF/ERK inhibition.

### Developmental time-dependent sensitivity to FGF modulation

Lineage tracing experiments have shown that ICM cells are committed to either Epi or PrE cell fate ^3^. Several studies have addressed the key role of FGF signaling in the control of Epi/PrE cell specification ^11-13^. However, it is not entirely clear how uncommitted ICM cells (NANOG and GATA6-positive cells) behave upon FGF modulation during this specification process particularly whether all or a subset) of uncommitted ICM cells can respond at any time to FGF modulation. We proposed from our data that the competency of uncommitted ICM cells to respond to FGF signaling is rapidly evolving with time. Indeed, we have identified four distinct periods where ICM cells exhibit distinct behaviors in response to FGF modulation (Fig. 6). From E2.75 to E3.25, ICM cells are sensitive to exogenous FGF stimulation as previously described ^6,11^. This suggests that while ICM cells exhibit heterogeneous levels of FGF transduction machinery *(Fgf4/Fgfr2)* ^18,19^, all of them can respond to increased levels of FGF activity. Consistent with previous studies on E3.25 *Fgf4* mutants ^8,9,19^, ICM cells are not affected by FGF/ERK inhibition during that period indicating that NANOG and GATA6 levels in ICM cells are not tightly regulated by FGF signaling at this stage. This period is followed by a stage (from E3.25 to E3.5) where ICM cells are still insensitive to FGF/ERK inhibition but lost their ability to respond to exogenous FGF stimulation. The reason for this is currently unclear. Whether this is due to dynamic regulation of components of the signaling cascade or intrinsic variations of the cell ability to respond to stimulation such as the phase of the cell cycle will require further investigations. From E3.5 to E3.75, ICM cells are highly sensitive to FGF/ERK inhibition as demonstrated by our observation that a short 4h treatment is sufficient to drive almost all ICM cells towards Epi cell fate (Fig. 4). However in the same period, ICM cells poorly respond to FGF stimulation. From E3.75, their ability to respond to modulation of FGF activity is progressively lost.

The mechanisms underlying the acquisition of such competence are currently unknown. One possibility could be that ICM cells with different sensitivity to FGF/ERK inhibition coexist within the ICM and are specified with different timing. Responsive ICM cells would be bias towards PrE fate and because they specify late, FGF/ERK inhibition at early timing would have no immediate impact on ICM composition. It has been proposed that inner cells generated during the second wave of asymmetric division express higher levels of *Fgfr2* and are biased towards PrE fate ^20,21^. However, whether these cells are specified later than inner cells generated from the first wave has not yet been determined. Another possibility would be that ICM will first give rise to Epi progenitors, as predicted by a mathematical modeling of ICM differentiation and confirmed *in situ* ^6^, that in turn would modify the remaining ICM cells and make them responsive to FGF/ERK signaling. The accumulation of FGF4 which is secreted by Epi cells is likely to mediate this switch as previously proposed ^22^. Above a certain threshold, which would be attained only when most Epi cells are specified, high FGF4 levels would trigger robust activation of FGF/ERK in remaining ICM cells and drive them towards PrE differentiation.

### Cell fate plasticity within Epi and PrE progenitor populations

Lineage tracing and chimera experiments have suggested that Epi progenitors exhibit less plasticity than PrE precursors ^12,13,23^. Our data also argue in favor of a model whereby, as they emerge, Epi progenitors loose their ability to be converted into PrE upon FGF stimulation. Indeed, embryos treated with FGF4 from E3.25 to E3.75 showed a significant bias towards PrE fate at the expense of Epi (Fig. 1B). While the number of NANOG-only ICM cells was reduced compared to control (9.8±0.8 vs. 5.2±1.0 NANOG-positive cells in control vs. treated embryos, Fig. 1Ba-b), it remains closer to the number of NANOG-only ICM cells present in E3.25 embryos before treatment (2.7±0.5 NANOG-positive cells, Fig. 1Aa). This suggests that FGF4 treatment prevented unspecified ICM cells to adopt Epi identity rather than affected the fate of already specified Epi progenitors. Between E3.5 and E3.75, FGF4 treatment had a mild impact on ICM composition so that 7.9±1.1 ICM cells remained Epi progenitors (compared to 9.8±0.8 cells in control conditions). Lastly, by E3.75, FGF4 treatment does not affect the composition of the ICM indicating that NANOG-positive cells are unlikely to be converted into PrE progenitors upon exogenous FGF stimulation.

Conversely, the ability of PrE progenitors to be converted into Epi in response to FGF inhibition seems to be modulated in time. This is exemplified by our observation that inhibition of FGF/ERK from E3.5 to E3.75 leads to an ICM almost exclusively composed of Epi progenitors (Fig. 4) while similar treatment has a less profound effect when applied from E3.75 to E4.5 (Fig. 1C).

Taken together our data fit with our current model of ICM cell specification and cellular plasticity and future studies using live imaging to address the temporal dynamics of these single and double-positive populations will undoubtedly give precious insights into these questions.

### Regulation of ICM cell specification by transcription and proteasome degradation

While several studies have addressed the interplay between NANOG, GATA6 and FGF signaling, it is still not clear how GATA6/NANOG positive uncommitted ICM cells mature into single positive cells and how FGF/ERK inhibition impacts on this process so that almost all cells are NANOG positive only. Our data indicate that inhibition of proteasome-mediated proteolysis or transcription) has a low impact on the establishment of GATA6/NANOG mutually exclusive expression from co-expressing ICM cells, while it severely impaired the effect of FGF/ERK inhibition.

Unexpectedly, inhibition of the proteasome led to a strong reduction of NANOG levels in Epi progenitors. This is surprising since NANOG protein stability in pluripotent stem cells has been shown to be tightly regulated through its PEST domain ^24^, ERK-mediated phosphorylation ^25,26^, interaction with stabilizing proteins such as the COP9 signalosome member COPS2 ^27^ or the deubiquitinase USP21 ^28^. Accordingly, MG132 treatment in ES cells leads to increased NANOG levels ^24,26,27,29^. Thus, our data reveal that NANOG levels are differentially regulated between ICM and pluripotent stem cells and point to additional mechanisms controlling NANOG stability *in vivo* during ICM cell specification. A likely possibility would be that an inhibitor of NANOG expression or stability is present in ICM cells and actively degraded by the proteasome between E3.5 and E3.75. It has been proposed that, among others ^30,31^, GATA6 is an important regulator of NANOG ^32-34^. Further experiments will be required to test its implication *in vivo* in ICM cells.

Although less pronounced than for NANOG, a positive regulation of GATA6 levels by the proteasome was observed in PrE progenitor. It would be interesting to determine the mechanisms regulating GATA6 stability and to test possible candidates such as interaction with the polycomb member BMI1 ^35^, destabilization of *Gata6* mRNA by UNR ^36^ or cAMP-dependent proteolysis that have been previously shown to regulate GATA6 levels in other contexts ^37,38^. A large number of evidences have pointed out the role of proteasome activity in stem and progenitor cells ^39^. In addition, other positive regulation on protein levels by the proteasome has been documented in various contexts, including through factors regulating the translation and the stability of mRNA during oocyte maturation ^40^. In general, a high dynamic regulation of the protein stability may be important to rapidly modulate cell fate decision in response to external cues.

Our data also show that contrary to NANOG levels in Epi, maintenance of high levels of GATA6 in PrE requires active transcription during the period when these progenitors are responsive to FGF/ERK inhibition (Fig 5D–E). Importantly, while inhibition of FGF/ERK signaling caused the complete downregulation of GATA6 in PrE progenitors already present in the embryos at the onset of the treatment, this was no longer the case when transcription was blocked. Similarly, blocking proteasome degradation prevent the conversion of ICM into Epi cell lineage upon FGF/ERK inhibition. Thus, our data show that the regulation of GATA6 levels by FGF signaling in PrE progenitors by the time of specification is manifold and acts at several levels including transcription and protein stability.

Taken together, our data shed light on the mechanisms involved in EPI/PrE specification which for the first time address the roles of transcription and proteasome.

## Material and methods

### Embryo collection and culture

Embryos from CD1 (Charles River laboratories, France) intercrosses were recovered at E2.75 and at E3.25 (expanding blastocyst) by flushing oviducts or uteri in M2 (Sigma Aldrich), washed twice and cultured inside their intact zona pellucida in 400μL KSOM (Millipore) in Nunc 4-well plates until the stages of interest at 37°C, 5% CO2. For treatment, we used FGF receptor inhibitor PD173074 (100nM, Sigma Aldrich), MEK inhibitor PD0325901 (1μM, Sigma Aldrich), flavopiridol (1μM, Selleckchem), MG132 (10μM, Sigma Aldrich) and FGF4 (1p.g/ml, R&D systems) supplemented with heparin (1p.g/ml, Sigma Aldrich). Experiments shown in Figure 2 (A–C) and Figure 3 were performed in parallel using same batches of embryos. All experiments were conducted according to the French and European regulations on care and protection of laboratory animals (EC Directive 86/609, French Law 2001-486 issued on June 6, 2001) and were approved by the Institut Pasteur ethics committee (n&#xB0; 2012-0011).

### Immunofluorescence

Embryos were fixed in 4&#x0025; PFA overnight at 4&°C and then washed three times in PBT (PBS, 0.1&% Triton X-100). Embryos were preincubated in blocking solution (PBT, 10&#x0025; FBS) during 1 hour and then incubated overnight at 4&°C. In this study, anti-GATA6 (1/100, AF1700, R&D Systems), anti-CDX2 (1/100, MU392A-UC, Biogenex), anti-NANOG (1/100, 8822, Cell Signaling; 1/100, 14-5761, eBioscience), and anti-SOX17 (1/100, AF1924, R&D systems) were used. After several washes in PBT, embryos were incubated with secondary antibodies coupled with Alexa 488nm, 546nm, 647nm (1/300, Invitrogen) and Hoechst (1/1000, Sigma Aldrich) 2 hours at room temperature and then washed in PBT. We manually counted the number of cells positive only for GATA6/SOX17/NANOG or double positive (GATA6 or SOX17/NANOG). Embryos were and analyzed using a SP5 (Leica) or LSM800 (Zeiss) confocal microscope. Images were acquired using glass bottomed microwell dishes (MatTek Corporation) using the same objective (Plan-apochromat 20x/NA 0.75), speed (700 Hz), pinhole 1 airy unit, and laser intensities with optical section thickness of 1.2-1.4μm. 16-bit resulting images were analyzed and eventually processed using Icy, Fiji and Photoshop CS5 softwares.

### Fluorescence quantification

3D stacks were analyzed using Icy software ^41^ (http://icy.bioimageanalysis.org), for which we designed a custom semi-automated analysis protocol available here. After selecting the nuclei staining channel, the protocol applies a wavelet decomposition of the 3D image volume to segregate image structures based on size and intensity. After optimizing the selection of wavelet scales and their respective thresholds, the protocol extracts one or more 3D regions of interest (ROI) for each nucleus. The selection of scales and thresholds is adapted to each embryo, depending on the signal-to-noise ratio. Finally, the protocol super-imposes the extracted ROI on the original data for visual inspection. In few instances, multiple ROI are detected inside the same nucleus, and are merged manually post-analysis. The mean of fluorescence intensity for GATA6 and NANOG channels for each ROI is then calculated.

### Blastocyst gene expression profile

mRNAs of 5 pooled blastocysts were isolated using RNeasy Plus Micro kit (Qiagen) according manufacturer instructions, reverse transcribed using Superscript VILO (Thermo Fischer Scientific) and analyzed using SYBR Green PCR Master Mix (Applied Biosystems). Mouse RT-qPCR primers were designed using Primer Blast (NCBI): Rps17-F 5'-CGCCATTATCCCCAGCAAGA-3&#x2019;, Rps17-R 5’ GCTGAGACCTCAGGAACGTAG-3&#x2019;, Pre-mRNA-F 5&#x2019;-TTTGTTGGTGTGTCCTGGCT-3&#x2019; and Pre-mRNA-R 5&#x2019;-CCACCCGGCTAATGAACACT-3&#x2019;.

### Statistical analysis

Graphs and statistical tests (t-test and Mann-Whitney) were performed using Prism (Graphpad).

## Acknowledgments

Imaging was performed for most parts at the Imagopole of Institut.

## Author contributions

Conceptualization: J.A., S.B., and M.C.-T.; Methodology: J.A., S.B., and M.C.-T.; Software: A.D.; Investigation: S.B., J.A., S.V.-P. and S.C; Writing-Original Draft: S.B., J.A. and M.C.-T; Writing-Review & editing: S.B., J.A. and M.C.-T; Supervision: J.A. and M.C.-T.; Funding acquisition: J.A. and M.C.-T.

---

**Supplementary Figure S1.**
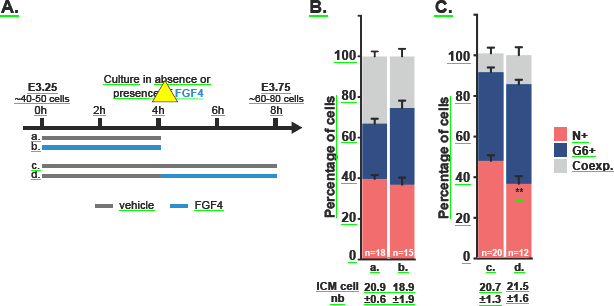
(related to Figures 3 and 4). Effect on ICM specification of exogenous FGF 4 stimulation between E3.25 and E3.75. (A)Schematic of the time schedule of vehicle (grey) and FGF4 (blue) treatment. (B-C) Distribution of ICM cells expressing NANOG (N+, red), GATA6 (G6+, blue) or both markers (Coexp., grey) in embryos cultured in presence or absence of FGF4. Control embryos plotted in Figure S2B are similar to Figure 1B. Error bars indicate SEM. n, number of embryos analyzed. Statistical Mann–Whitney tests are indicated when significant (**, p < 0.01).

**Supplementary Figure S2.**
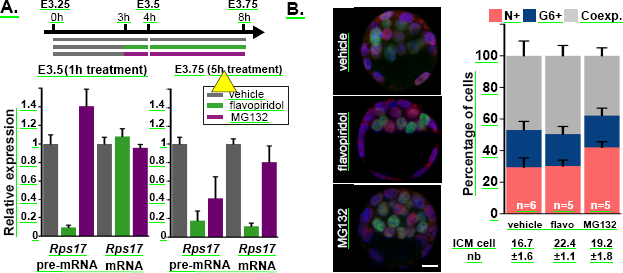
(related to Figure 5). Effect of modulating transcription/proteasome activity during ICM specification. (A) Schematic of the time schedule of inhibitor treatment. Green, purple and grey lines indicate the culture periods in the presence of flavopyridol, MG132 and DMSO (vehicle), respectively. RT-qPCR expression analysis of Rps17 pre-mRNA
and Rps17 mRNA in embryos cultured 1h and 5h with/without the drugs. (B) Immunodetection of NANOG (green) and GATA6 (red) in embryos cultured 1h (from 3h to 4h) in the presence/absence of drugs. Pictures correspond to a projection of 5 confocal optical slices. Scale bar: 20μm. Distribution of ICM cells expressing NANOG (N+, red), GATA6 (G6+, blue) or both markers (Coexp., grey) in embryos cultured for the indicated period of times. Error bars indicate SEM. n, number of embryos analyzed.

